# The DNA repair associated protein Gadd45*γ* regulates the temporal coding of immediate early gene expression and is required for the consolidation of associative fear memory

**DOI:** 10.1101/265355

**Authors:** Xiang Li, Paul R. Marshall, Laura J. Leighton, Esmi L. Zajaczkowski, Timothy W. Bredy, Wei Wei

**Affiliations:** Cognitive Neuroepigenetics Laboratory, Queensland Brain Institute, The University of Queensland, QLD 4072, Australia

## Abstract

**Visual abstract.**
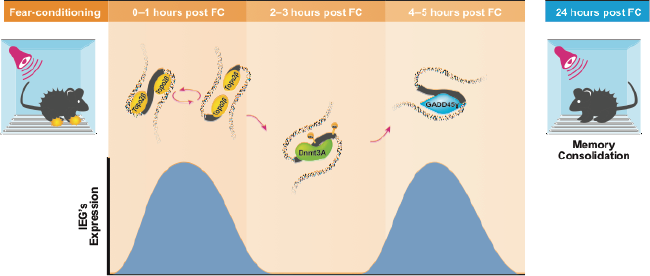

We have identified a member of the Growth arrest and DNA damage (Gadd45) family, Gadd45*γ*, which is known to be involved in the regulation of DNA repair, as a key player in the formation of associative fear memory. Gadd45*γ* regulates the temporal dynamics of learning-induced immediate early gene (IEG) expression in the prelimbic prefrontal cortex through its interaction with DNA double-strand break (DSB)-mediated changes in DNA methylation. Our findings suggest a two-hit model of experience-dependent IEG activity and learning that comprises 1) a first wave of IEG expression governed by DSBs followed by an increase in DNA methylation, and 2) a second wave of IEG expression associated with Gadd45*γ* and active DNA demethylation at the same site, which is necessary for memory consolidation.

**Significance statement:** How does the pattern of immediate early gene (IEG) transcription in the brain relate to the storage and accession of information, and what controls these patterns? This paper explores how GADD45γ, a gene that is known to be involved with DNA modification and repair, regulates the temporal coding of IEGs underlying associative learning and memory. We reveal that, during fear learning, GADD45*γ* serves to act as a coordinator of IEG expression and subsequent memory consolidation by directing temporally specific changes in active DNA demethylation at the promoter of plasticity-related IEGs.

## Introduction

Memory consolidation has been shown to require learning-induced changes in immediate early gene (IEG) expression, protein synthesis, and neuronal structural changes (1, 2). Building on this foundation, recent work has demonstrated that there is yet another layer of regulatory control over experience-dependent gene expression and memory, which involves epigenetic processes such as histone and DNA modification as well as the coordination of such processes by various classes of noncoding RNA (3, 4). With respect to DNA modification, our appreciation of the role of this epigenetic mechanism in learning and memory has increased dramatically with the discovery of a role for both DNA methylation- and active demethylation-related changes in gene expression and memory formation (5, 6).

Several active DNA demethylation pathways have been proposed, and each has been shown to be involved in regulating gene expression related to plasticity and memory (7). The first involves hydroxylation of 5-methylcytosine (5-mC) by Tet1-3, followed by further oxidation to form 5-formylcytosine and then 5-carboxylcytosine, which is removed by DNA glycosylases (TDG and MBD4) through base excision repair. We and others have recently shown that this pathway is associated with memory formation (5, 8, 9). The second pathway involves deamination of 5-hmC by AID to form 5-hydroxymethyluridine, which is then removed by TDG/MBD4-mediated base excision repair, which has also been demonstrated to play a role in activity-induced gene expression (10). Finally, the third, and perhaps most direct pathway involves members of the Gadd45 protein family, which remove 5-mC by nucleotide excision repair. Gadd45α is required for active DNA demethylation as it functions in a complex that includes other DNA repair enzymes (11-13). Furthermore, knockdown of Gadd45β has significant effects on learning, although reports differ with regards to whether this knockdown leads to enhancement (14) or impairment of memory (15).

Emerging evidence indicates that DNA double strand breaks (DSB) and DNA repair may also be required for the gene expression that underlies memory formation (16, 17). These processes interact with dynamic changes in DNA methylation and can trigger methylation iteself (18-22). In line with this, together with its potential role in active DNA demethylation, GADD45β is recruited to genomic loci in response to genotoxic stimuli that generate DSB’s (e.g. radiation) (23, 24). Based on these observations, we questioned whether members of the Gadd45 family influence gene expression and memory by interacting with DSBs, DNA repair and DNA methylation. Our results indicate that several plasticity-related IEGs, including activity-regulated cytoskeleton-associated protein (Arc), fos proto-oncogene (Fos),neuronal PAS domain protein 4 (Npas4) and a newly identified IEG, cysteine-rich angiogenic inducer 61 (Cyr61), are subject to DSBs upon their induction, which is followed by a rapid increase in DNA methylation and a time-dependent recruitment of DNA binding proteins. Surprisingly, the mRNA levels of these IEGs peak twice in response to cued fear learning, with markers of DSBs *γ*H2A.X and topoisomerase 2-beta (Topo ⃦β) corresponding to the first peak and Gadd45*γ*-mediated DNA repair regulating the second peak. In addition, we have found that knockdown of the Gadd45*γ* target Cyr61 also impairs the consolidation of fear memory.

## Results

### Learning-induced Gadd45*γ* expression in the prelimbic prefrontal cortex is required for the formation of cued fear memory

To establish which members of the Gadd45 family are involved in regulating gene expression in the prelimbic prefrontal cortex (PLPFC) during fear learning, we examined Gadd45α, Gadd45β and Gadd45*γ* mRNA expression following stimulation of cultured neurons with potassium chloride, and following learning in adult mice. Contrary to previous findings in the striatum, hippocampus and amygdala (14, 15) only Gadd45γ mRNA showed a significant increase in response to cued fear conditioning (Fig. 1A-C). In primary cortical neurons *in vitro,* similar to previous observations, neural activation led to a significant increase in Gadd45β and Gadd45*γ*, but not Gadd45α, mRNA transcript levels (SI Appendix Fig. S1A-C.). However, the level of mRNA expression does not necessarily reflect the functional relevance of a given gene (25). Given previous findings of a role for Gadd45β in regulating contextual fear memory, we therefore designed shRNAs against all three members of the Gadd45 family according to established protocols (26), and validated these *in vitro* (SI Appendix Fig. S1D-H). Each shRNA was then separately infused into the PLPFC (SI Appendix Fig. S2A-C) at least 1 week prior to behavioural training (Fig. 1D). There was no effect of knockdown of any member of the Gadd45 family on the acquisition of freezing behaviour during cued fear learning (Fig. 1E). Knockdown of Gadd45α and Gadd45β had no effect on memory retention. In contrast, Gadd45*γ* shRNA-treated mice showed a significant impairment in fear memory (Fig. 1F-H). Furthermore, there was no significant difference between control and Gadd45*γ* shRNA-treated mice on locomotor or anxiety-like behaviour in the open field test (SI Appendix Fig. S2D-F.). Together, these data demonstrate a critical role for Gadd45*γ*, but not Gadd45*α* or Gadd45*β*, within the PLPFC in the regulation of cued fear memory.

**Fig. 1.**
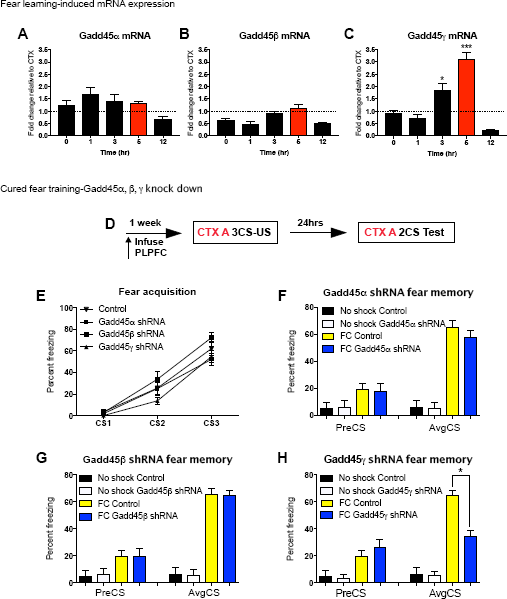
Gadd45*γ* expression is activity-dependent and necessary for cued fear learning in the PLPFC. Error bars represent S.E.M. *p < 0.05 **p < 0.01 *** p < 0.001. Figure (A-C) shows fold change for fear conditioned animals which is calculated relative to its own context control at that time point. Dotted line represents average of context controls. (A) Following cued fear conditioning there was no significant change in Gadd45α or (B) Gadd45*γ* mRNA levels. (C) Although there was a significant increase in Gadd45*γ* 3 and 5 hours postconditioning (n = 4 per group; F(9,30) = 36.33 p <0.001;*Sidak post-hoc analysis compared CTX 3hr to FC 3hr and CTX 5hr to FC 5hr) (D) Time course of behavioural training and shRNA infusion. (E) There were no significant differences between groups on percent freezing during fear acquisition (F) There were also no significant differences of percent freezing during the fear test for animals infused with either Gadd45α shRNA (G) or Gadd45β shRNA lenti-virus (H) when compared to control. However, there was a significant decrease in the percent freezing for animals infused with Gadd45*γ* shRNA compared to control at test (n = 14 per group; F(3, 24) = 27.37 p <0.001; *Sidak-post hoc analysis, FC Control vs. FC Gadd45*γ* shRNA). FC: fear conditioned; CTX: context exposure control; CS: conditioned stimulus (tone); US: unconditioned stimulus (shock).

### Fear learning leads to a distinct pattern of IEG expression in the PLPFC

We next assumed a candidate gene approach to obtain a detailed understanding of the mechanism by which Gadd45*γ* influences the formation of fear memory. The IEGs Arc, Fos, and Npas4 represent prime choices because of their well-known role in regulating fear-related learning and memory (27-30). In addition, lEGs have been shown to be rapidly induced by DSBs, which are later subject to repair (17). We also included the newly discovered IEG, Cyr61, as it is also shown to be expressed in the brain and is induced by neural activity (31, 32). An initial analysis of IEG mRNA levels revealed that Arc, Fos, Npas4 and Cyr61 exhibit two significant peaks of expression in the PLPFC in response to cued fear conditioning (Fig. 2A-D). This is reminiscent of earlier observations by Izquierdo and colleagues in which two waves of transcription, one occurring immediately after training and one occurring 3-6 hours later, were shown to be required for the formation of hippocampal-dependent fear memory (33), as well as a variety of other reports in which double IEG peaks have been observed in the context of learning (28, 34).

**Fig. 2.**
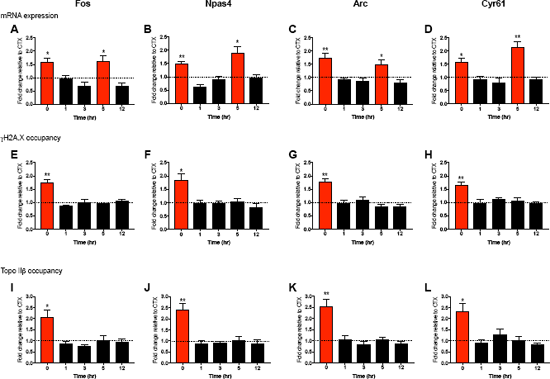
Fear learning leads to a distinct pattern of IEG expression in the PLPFC regulated by DSBs. All panels show fold change for fear conditioned animals which is calculated relative to its own context control at that time point. Dotted line represents average of context controls. Error bars represent S.E.M. *p < 0.05 **p < 0.01 *** p < 0.001. (A) There was a significant increase in mRNA 0 hours and 5 hours post fear conditioning for Fos (n = 6 per group; F(9,41) = 4.20 p <0.001; *Sidak post-hoc analysis comparing CTX 0hr to FC 0hr and CTX 5hr to FC 5hr) (B) Npas4 (n = 6 per group; F(9,40) = 7.04 p <0.001), (C) Arc (n = 6 per group; F(9,40) = 6.42 p <0.001 and (D) Cyr61 (n = 6 per group; F(9,42) = 8.67 p <0.001). (E) There was also a significant increase in *y*H2A.X occupancy immediately following cued fear conditioning for Fos (n = 6 per group; F(9,40) = 4.16 p <0.001; Sidak post-hoc analysis comparing CTX 0hr to FC 0hr), (F) Npas4 (n = 6 per group; F(9,40) = 2.95 p <0.001), (G) Arc (n = 6 per group; F(9,40) = 4.87 p <0.001) and (H) Cyr61 (n = 6 per group; F(9,40) = 2.20 p <0.05). (E) We also observed a significant increase in topoisomerase IIβ binding immediately following cued fear conditioning for Fos (n = 6 per group; F(9,40) = 4.45 p <0.001; *Sidak post-hoc analysis comparing CTX 0hr to FC 0hr), (F) Npas4 (n = 6 per group; F(9,40) = 8.82 p <0.001), (G) Arc (n = 6 per group; F(9,40) = 7.83 p <0.001) and (H) Cyr61 (n = 6 per group; F(9,40) = 4.06 p <0.001).

### IEG activity is regulated by DSBs and time-dependent increases in DNA methylation

It has been observed *in vitro* that Fos and Arc require DSBs for their activation (17), and that DSBs lead to the recruitment of the Gadd45 family of repair enzymes (22). To determine whether these IEGs are subject to DSBs in the adult brain, we probed their proximal promoters for evidence of learning-induced DSBs following fear conditioning. γH2A.X has been shown to be an excellent marker for DSBs (35), and Topo IIβ is known to be involved in the repair of DSBs (36). All four IEGs exhibited a significant increase in both γH2A.X and Topo IIβ binding immediately following fear conditioning, the same time at which the first peak of gene expression for all four IEGs occurred (Fig. 2E-L). However, this did not explain the origin of the second peak of gene expression. It has been previously shown that DSBs induced in neuons by learning are quickly repaired by Topo IIB [ref] and our data also supported this finding. Given that DSBs are repaired before the second wave of transcription at these loci, we considered other mechanisms that could maintain the locus in a poised state following the repair of the originating DSB. DNA methylation has been shown to increase at sites of DSBs (22) and increased DNA methylation is associated with memory formation (37), so we investigated the methylation state at these loci. DNMT3A chromatin immunoprecipitation (ChIP) and methylated DNA immunoprecipitation (MeDIP) revealed a significant increase in DNMT3A binding at the site of DSB in all IEGs (Fig 3A-D), which was followed by an increase in 5-mC up to 3 hours post fear conditioning (Fig 3E-H).

**Fig. 3.**
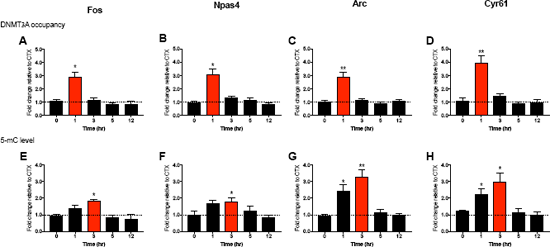
Fear conditioning associated with time-dependent increases in DNA methylation. All figures show fold change for fear conditioned animals which is calculated relative to its own context control at that time point. Dotted line represents average of context controls. Error bars represent S.E.M. *p < 0.05 **p < 0.01 *** p < 0.001. (A) A significant increase in DNMT3A occupancy was observed 1hr after fear conditioning for Fos (n = 6 per group; F(9,40) = 7.77 p <0.001; * Sidak post-hoc analysis compared Context 1hr to FC 1hr), (B) Npas4 (n = 6 per group; F(9,40) = 11.40 p <0.001), (C) Arc (n = 6 per group; F(9,40) = 10.20 p <0.001) and (D) Cyr61 (n = 6 per group; F(9,40) = 15.30 p <0.001). (E) A significant increase in 5-mC levels as measured by methylated DNA immunoprecipitation was found post fear conditioning for Fos (n = 6 per group; F(9,40) = 2.67 p <0.001; *Sidak post-hoc analysis compared CTX 1hr to FC 1hr and CTX 3hr to FC 3hr), (F) Npas4 (n = 6 per group; F(9,40) = 2.16 p <0.05), (G) Arc (n = 6 per group; F(9,40) = 10.52 p <0.001) and (H) Cyr61 (n = 6 per group; F(9,40) = 5.52 p <0.001).

### Gadd45*γ* regulates learning-induced IEG expression in a temporally specific manner through interaction with DNA methylation

Next, to determine whether Gadd45*γ* was targeting the DSBs, DNA methylation, or both, we performed ChIP for Gadd45*γ* occupancy on the tissue derived from fear-conditioned animals. There was a significant increase in Gadd45γ binding at the 5 hour time point for all IEGs (Fig. 4A-D). Subsequent control experiments determined that this binding was specific to Gadd45*γ* as ChIP for Gadd45α or Gadd45β revealed no binding at the same loci (SI Appendix Fig. S3A-B.). Additionally, there was no significant binding of Gadd45*γ* at distal promoter regions of Cyr61 (SI Appendix Fig. S3C). Importantly, Gadd45*γ* knockdown significantly reduced the presence of Gadd45*γ* (Fig. 4E-H) and blocked mRNA expression at the second peak only (Fig. 4I-L). In addition, 5-mC levels declined sharply at the time point at which Gadd45*γ* bound with control shRNA on board, whereas Gadd45*γ* knockdown led to persistently high levels of 5-mC (Fig. 4M-P). Together, these data suggest that, although DSBs may be required for the initial activation of the IEG expression, the second critical peak of IEG expression during the consolidation phase of memory is regulated by Gadd45*γ* mediated changes in DNA methylation.

**Fig. 4.**
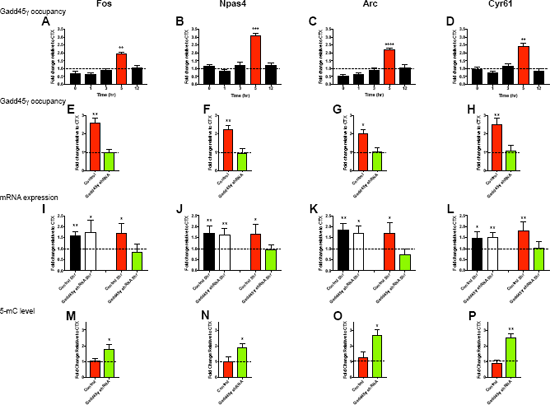
Gadd45*γ* regulates learning-induced IEG expression in a temporally specific manner through interaction with DNA methylation. All figures show fold change for fear conditioned animals which is calculated relative to its own context control at that time point. Dotted line represents average of context controls. Error bars represent S.E.M. *p < 0.05 **p < 0.01 *** p < 0.001. There was a significant increase in Gadd45*γ* occupancy at 5hr for (A) Fos (n = 6 per group; F(9,40) = 7.33 p <0.001; *Sidak post-hoc analysis compared CTX 5hr to FC 5hr), (B) Npas4 (n = 6 per group; F(9,40) = 17.02 p <0.001), (C) Arc (n = 6 per group; F(9,40) = 8.51 p <0.001) and (D) Cyr61 (n = 6 per group; F(9,40) = 11.54 p <0.001). Gadd45*γ* shRNA led to significantly less occupancy for (E) Fos (n = 4-6 per group; t(10) = 5.02, p <0.001), (F) Npas4 (t(10) = 3.88, p <0.001), (G) Arc (t(10) = 2.93, p <0.001) and (H) Cyr61 (t(10) = 3.35, p <0.001). Gadd45*γ* knockdown also led to a significant decrease in mRNA at 5hr for (I) Fos (n = 6 per group; F(3,16) = 5.77 p<0.001; *Sidak post-hoc analysis comparing Control 5hr to Gadd45*γ* shRNA 5hr), (J) Npas4 (F(3,16) = 5.82 p <0.001), (K) Arc (F(3,16) = 10.65 p <0.001) and (L) Cyr61 F(3,16) = 5.63 p <0.001). Gadd45*γ* shRNA led to significantly more methylation at 5 hours compared to Control for (M) Fos (n = 4-6 per group; t(7) = 2.02, p <0.05), (F) Npas4 (t(7) = 2.15, p <0.05), (G) Arc (t(10) = 2.69, p <0.05) and (H) Cyr61 (t(10) = 4.96, p <0.001).

### Knockdown of Cyr61 impairs the formation of fear memory

Many of these targets of Gadd45*γ* have previously been shown to influence the formation of fear memory, including Fos (28), Npas4) (30), and Arc (29). In order to extend these findings, we designed and validated an shRNA against the newly identifed IEG, Cyr61 (SI Appendix Fig. S4A-B). We then infused this lenti-viral shRNA into the PLPFC following the same behavioural timeline as Gadd45*γ* knockdown (Fig. 1D and Fig. 5A). Knockdown of Cyr61 had no significant effect on the acquisition of freezing during cued fear training and no effect on locomotor activity or anxiety-like behaviour in the open field test. However, knockdown of Cyr61 led to a significant impairment in fear memory, although notably to a much lesser degree than Gadd45γ (Fig. 5B-F).

**Fig. 5.**
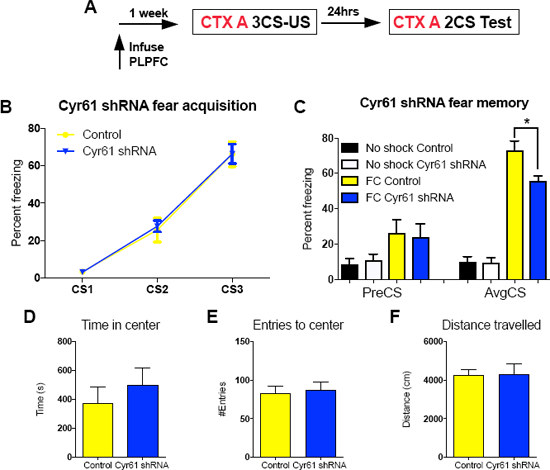
The immediate early gene Cyr61 is necessary for cued fear learning in the PLPFC. Error bars represent S.E.M. *p < 0.05 **p < 0.01 *** p < 0.001. (A) Time course of behavioural training and shRNA infusion. (B) There were no significant differences between the Cyr61 shRNA and control groups on percent freezing during fear acquisition. (C) There was a significant decrease in the percent freezing for animals infused with Cyr61 shRNA compared to control at test (n = 7 per group; F(3, 24) = 27.37 p <0.001; *Sidak post-hoc analysis FC Control vs. FC Cyr61 shRNA). During open field testing there were no significant differences between control and Cyr61 shRNA animals for (D) time spent in the center, (E) entries into the center or (F) distance traveled.

## Discussion

We have discovered that there is a tight temporal relationship between learning-induced DSBs, DNA repair and DNA (de)methylation in the regulation of learning-induced IEG expression, and highlight Gadd45*γ* as a central regulator of the temporal coding of IEG transcription that is required for fear memory consolidation. To our knowledge this is the first demonstration of a casual effect of Gadd45*γ* in learning, and the first model synthesising the functional interaction of DSBs, DNA methylation and DNA repair mediated DNA demethylation in a learning context. This model is significant as it provides a testable platform for further experimentation. Additionally, the interpretation of the data adds mechanistically to the literature describing memory consolidation as a process which is not simply linear and proportional to the passage of time, but is instead continually and dynamically stabilized and destabilized across time.

Whereas previous studies observed that global Gadd45β knockout affected contextual but not cued fear conditioning (15) we have found that Gadd45γ; but not Gadd45α or Gadd45β, is required in the PLPFC for cued fear conditioning (Fig. 1A-H). This suggests that these genes may in fact be region-specific, and thus also task-specific. Complementary to this is the observation of potential selectivity of Gadd45*γ* for IEGs. This is supported by the data showing that all of the IEGs selected in this study are bound at the same 5 hour time point by Gadd45*γ*, with similarly reduced mRNA expression at this time point following knockdown. This targeting of IEGs is also supported by the observation that knockdown of Gadd45*γ*, which binds a variety of IEGs, impairs fear conditioning by a considerably larger margin than the single IEG knockdown of Cyr61. This selectivity of IEGs is interesting because it has been shown previously that Gadd45β binds more slowly to transcribed neurotrophic factors (13). Thus, in addition to specificity for different brain regions and potential cell types, members of the Gadd45 family may also be targeting different classes of genes altogether.

Perhaps even more intriguing is the fact that while IEGs are bound by Gadd45*γ*, these IEGs are also a hot spot for DNA breaks and dynamic DNA methylation. Our time-course analysis revealed that the DSBs that occur in response to learning are followed by an increase in DNA methylation (Fig. 3A-H). Until now this change in DNA methylation following DSBs has been suggested to occur in only a few cells after aberrant repair (22); our data showing that Gadd45*γ*-mediated demethylation controls the second peak of IEG expression in the PLPFC suggests instead that this phenomenon may be widespread in the brain and used functionally for priming (20-22). Further validating this interpretation is the recent work that has shown that both 5-formylcytosine, and 5-mC within CA dinucleotides, can serve to epigenetically prime gene activity throughout the brain (38, 39).

Counterintuitively, Gadd45*γ* is targeting these IEGs, but only at the 5 hour timepoint corresponding to the second wave of IEG expression (Fig 4A-L). This observation is made more intriguing by the long-held position that there are two time periods during which protein synthesis inhibitors impair consolidation, with predictions of these two critical windows being immediately following learning and 3-6 hours later (40-42). The dual observation that these genes are subject to both DSBs and Gadd45*γ* targeting is not to be ignored as it suggests a two-tier process whereby DSBs are needed to activate IEGs at the first time point, and activity-dependent DNA demethylation guided by Gadd45}γ is critical for this IEG pattern to trigger processes required for long-term memory at the second.

As described by Izquierdo and colleagues (33), the data on double peaks of gene expression fit within the lingering consolidation hypothesis, which states that there are processes which occur across time and result in continued destabilization and restabilization of memory traces, contrasted to the simpler model describing a linear relationship between memory strength and time (43). This distinction is theoretically important as the latter assumes certain time periods, such as 24 hours, could be used as a control where manipulations should have no effect. In fact, Katche *et al* (2010) have shown that manipulation of cFos 24hrs after initial training results in impairments of long-term memory storage. Here we add mechanistically to this model by suggesting that a DNA modification switch that is activated by DSBs and regulated by DNA repair is critical for this process.

The model would make further predictions that DNA demethylation may not only be critical for the initial learning-induced induction and encoding of information at the level of gene expression, but also indicate a retrieval-induced recapitulation of transcriptional activity. Supporting this idea is the recent observation that retrieval-induced Tet3 gene expression is necessary for retrieval and reconsolidation (44), as well as selective retrieval impairment by infusion of DNA methyltransferase inhibitors into the PLPFC (45), although further work is required to establish this. It also remains to be seen whether the phenomenon of DSBs followed by DNA methylation unmasking generalizes to other classes of genes or whether other mechanisms of DNA modification also follow this time course. Nonetheless, our data imply a two-hit model of IEG activation whereby the initial activation is dependent on DSBs, while the second wave that is critical for memory consolidation depends on Gadd45*γ*-mediated DNA demethylation and repair.

## Acknowledgements

The authors gratefully acknowledge grant support from the NIH (5R01MH105398-TWB) and the NHMRC (APP1033127-TWB). XL was supported by postgraduate scholarships from the University of Queensland and the ANZ trustees Queensland for medical research. PM is supported by postgraduate scholarships from NSERC and the University of Queensland. LL is supported by postgraduate scholarships from the Westpac Bicentennial Foundation and the University of Queensland. We would also like to thank Ms. Rowan Tweedale for helpful editing of the manuscript.

## References

1. Izquierdo I, McGaugh JL (2000) Behavioural pharmacology and its contribution to the molecular basis of memory consolidation. Behav Pharmacol 11(7-8):517–34.

2. Kandel ER (2001) The Molecular Biology of Memory Storage: A Dialogue Between Genes and Synapses. Science 294(5544):1030–1038.

3. Spadaro PA, Bredy TW (2012) Emerging role of non-coding RNA in neural plasticity, cognitive function, and neuro psychiatric disorders. Front Genet 3(JUL):1–16.

4. Baker-Andresen D, Ratnu VS, Bredy TW (2013) Dynamic DNA methylation: A prime candidate for genomic metaplasticity and behavioral adaptation. Trends Neurosci 36(1):3–13.

5. Li X, et al. (2014) Neocortical Tet3-mediated accumulation of 5-hydroxymethylcytosine promotes rapid behavioral adaptation. Proc Natl Acad Sci 111(19):7120–5.

6. Miller CA, et al. (2010) Cortical DNA methylation maintains remote memory. Nat Neurosci 13(6):664–666.

7. Li X, Wei W, Ratnu VS, Bredy TW (2013) On the potential role of active DNA demethylation in establishing epigenetic states associated with neural plasticity and memory. Neurobiol Learn Mem 105:125–132.

8. Kaas G, et al. (2013) TET1 controls CNS 5-Methylcytosine Hydroxylation, active DNA demethylation, gene transcription, and memory formation. Neuron 79(6):1086–1093.

9. Rudenko A, et al. (2013) Tet1 is critical for neuronal activity-regulated gene expression and memory extinction. Neuron 79(6):1109–1122.

10. Ratnu VS, Wei W, Bredy TW (2014) Activation-induced cytidine deaminase regulates activity-dependent BDNF expression in post-mitotic cortical neurons. Eur J Neurosci 40(7):3032–3039.

11. Rai K, et al. (2008) DNA Demethylation in Zebrafish Involves the Coupling of a Deaminase, a Glycosylase, and Gadd45. Cell 135(7):1201–1212.

12. Barreto G, et al. (2007) Gadd45a promotes epigenetic gene activation by repair-mediated DNA demethylation. Nature 445(7128):671–675.

13. Ma DK, Guo JU, Ming GL, Song H (2009) DNA excision repair proteins and Gadd45 as molecular players for active DNA demethylation. Cell Cycle 8(10):1526–1531.

14. Sultan FA, Wang J, Tront J, Liebermann DA, Sweatt JD (2012) Genetic Deletion of gadd45b, a Regulator of Active DNA Demethylation, Enhances Long-Term Memory and Synaptic Plasticity. J Neurosci 32(48):17059–17066.

15. Leach PT, et al. (2012) Gadd45b knockout mice exhibit selective deficits in hippocampus-dependent long-term memory. Learn Mem 19(8):319–324.

16. Su Y, Ming GL, Song H (2015) DNA damage and repair regulate neuronal gene expression. Cell Res 25(9):993–994.

17. Madabhushi R, et al. (2015) Activity-Induced DNA Breaks Govern the Expression of Neuronal Early-Response Genes. Cell 161(7):1592–1605.

18. Niehrs C, Schäfer A (2012) Active DNA demethylation by Gadd45 and DNA repair. Trends Cell Biol 22(4):220–227.

19. Pastink A, Eeken JCJ, Lohman PHM (2001) Genomic integrity and the repair of double-strand DNA breaks. Mutat Res – Fundam Mol Mech Mutagen 480-481:37–50.

20. Wossidlo M, et al. (2010) Dynamic link of DNA demethylation, DNA strand breaks and repair in mouse zygotes. EMBO J 29(11):1877–1888.

21. O’Hagan HM, Mohammad HP, Baylin SB (2008) Double strand breaks can initiate gene silencing and SIRT1-dependent onset of DNA methylation in an exogenous promoter CpG island. PLoS Genet 4(8):e1000155.

22. Morano A, et al. (2014) Targeted DNA methylation by homology-directed repair in mammalian cells. Transcription reshapes methylation on the repaired gene. Nucleic Acids Res 42(2):804–821.

23. Smith ML, et al. (1996) Antisense GADD45 expression results in decreased DNA repair and sensitizes cells to u.v.-irradiation or cisplatin. Oncogene 13(10):2255–63.

24. Gu X, et al. (2000) A DNA damage signal is required for p53 to activate gadd45. Cancer Res 60(6): 1711–1719.

25. Nainar S, Marshall PR, Tyler CR, Spitale RC, Bredy TW (2016) Evolving insights into RNA modifications and their functional diversity in the brain. Nat Neurosci 19(10):1292–8.

26. Lin Q, et al. (2011) The brain-specific microRNA miR-128b regulates the formation of fear-extinction memory. Nat Neurosci 14(9): 1115–1117.

27. Tischmeyer W, Grimm R (1999) Activation of immediate early genes and memory formation. Cell Mol Life Sci 55(4):564–574.

28. Katche C, et al. (2010) Delayed wave of c-Fos expression in the dorsal hippocampus involved specifically in persistence of long-term memory storage. Proc Natl Acad Sci 107(1):349–354.

29. Ploski JE, et al. (2008) The Activity-Regulated Cytoskeletal-Associated Protein (Arc/Arg3.1) Is Required for Memory Consolidation of Pavlovian Fear Conditioning in the Lateral Amygdala. J Neurosci 28(47):12383–12395.

30. Ploski JE, Monsey MS, Nguyen T, DiLeone RJ, Schafe GE (2011) The neuronal PAS domain protein 4 (Npas4) is required for new and reactivated fear memories. PLoS One 6(8):e23760.

31. Sakuma K, et al. (2015) Temporal and spatial transcriptional fingerprints by antipsychotic or propsychotic drugs in mouse brain. PLoS One 10(2).

32. Albrecht C, et al. (2000) Muscarinic acetylcholine receptors induce the expression of the immediate early growth regulatory gene CYR61. J Biol Chem 275(37):28929–36.

33. Igaz LM, Vianna MRM, Medina JH, Izquierdo I (2002) Two time periods of hippocampal mRNA synthesis are required for memory consolidation of fear-motivated learning. J Neurosci 22(15):6781–9.

34. Lefer D, Perisse E, Hourcade B, Sandoz J, Devaud J-M (2012) Two waves of transcription are required for long-term memory in the honeybee. Learn Mem 20(1):29–33.

35. Kuo LJ, Yang LX (2008) Gamma-H2AX – a novel biomarker for DNA double-strand breaks. In Vivo 22(3):305–309.

36. Calderwood SK (2016) A critical role for topoisomerase IIb and DNA double strand breaks in transcription. Transcription 7(3):75–83.

37. Day JJ, Sweatt JD (2010) DNA methylation and memory formation. Nat Neurosci 13(11):1319–1323.

38. Song CX, et al. (2013) Genome-wide profiling of 5-formylcytosine reveals its roles in epigenetic priming. Cell 153(3):678–691.

39. Stroud H, et al. (2017) Early-Life Gene Expression in Neurons Modulates Lasting Epigenetic States. Cell. doi:10.1016/j.cell.2017.09.047.

40. Grecksch G, Matthies H (1980) Two sensitive periods for the amnesic effect of anisomycin. Pharmacol Biochem Behav 12(5):663–665.

41. Freeman FM, Rose SP, Scholey a B (1995) Two time windows of anisomycin-induced amnesia for passive avoidance training in the day-old chick. Neurobiol Learn Mem 63(3):291–295.

42. Chew SJ, Vicario DS, Nottebohm F (1996) Quantal Duration of Auditory Memories. Science 274(5294):1909–1914.

43. Dudai Y, Eisenberg M (2004) Rites of passage of the engram: Reconsolidation and the lingering consolidation hypothesis. Neuron 44(1):93–100.

44. Liu C, et al. (2017) Retrieval-Induced Upregulation of Tet3 in Pyramidal Neurons of the Dorsal Hippocampus Mediates Cocaine-Associated Memory Reconsolidation. Int J Neuropsychopharmacol. pyx099.

45. Han J, et al. (2010) Effect of 5-aza-2-deoxycytidine microinjecting into hippocampus and prelimbic cortex on acquisition and retrieval of cocaine-induced place preference in C57BL/6 mice. Eur J Pharmacol 642(1-3):93–98.

